# An integrated system for comprehensive mouse peripheral vestibular function evaluation based on Vestibulo-ocular Reflex

**DOI:** 10.1101/2025.02.21.639406

**Authors:** Tong Zhao, Shijie Xiao, Wenda Liu, Jinhao Zhong, BinXian Sun, Fangyi Chen

**Author notes:** Corresponding author: Fangyi Chen. These authors contributed equally to this work (Tong Zhao, Shijie Xiao).

## Abstract

In the realms of both vestibular and auditory research, conducting vestibular function tests is essential. However, unlike the auditory function tests which utilize standard equipment such as the Auditory Brainstem Response (ABR) device, there is no equivalent widely adopted apparatus for vestibular tests. This is largely due to the intricate nature of the vestibular system and the challenges associated with assessing its functions. Vestibulo-ocular reflexes (VORs) are the compensatory ocular reflexes that ensure stable vision during head motion. VORs are widely used in clinics for diagnosing the vestibular deficit. In the research field, VORs, including angular VOR (aVOR) or off-vertical axis rotation (OVAR) tests, have been used by various groups to evaluate the mouse vestibular function. However, the effectiveness of VOR tests has not been systematically evaluated with proper animal models, and the lack of commercial equipment hampers its accessibility, confining vestibular testing to a select few labs. In this study, we developed an integrated instrument system with both aVOR and OVAR modes for evaluating mouse vestibular function. To demonstrate its efficacy, peripheral vestibular animal models, 1) Vestibulotoxicity drugs 3,3’-iminodiproprionitrile (IDPN, 2 mg/g and 4 mg/g) induced; 2) Critical MET-related mutant (*Cdh23^v2J/v2J^* and *TMC1^-/-^*); 3) Vestibulo-specific mutant (*Zpld1^-/-^* for semicircular canal dysfunction and *Otop1^tlt/tlt^* for otoconia deficient; 4) Unilateral vestibular lesion (UVL) model by injecting gentamicin into horizontal semicircular canal, were constructed and evaluated with the system. The results showed 1) Quantification of the vestibular deficit is achieved in a daily manner; 2) Both the otolith organ and semicircular canals can be assessed respectively; and 3) The lesion side of UVL can be identified. During an 8-week study of IDPN vestibulotoxicity, the vestibular function of 3 groups of 20 animals was evaluated at 15 test days. These test results reveal the potential of our system as a standard system for evaluating common vestibular deficits in mice.

## Introduction

The vestibular system is responsible for detecting changes in head and body position, controlling gaze, and facilitating higher-level cognitive perception functions(Mucha, Fedor, & DeMarco, 2018). It consists of three semicircular canals and two otolith organs (utricle and saccule), which mediate sensitivity to rotational and linear (including gravity) head movements, respectively(Lim & Brichta, 2016). In recent years, studies on the pathological mechanisms behind vestibular dysfunction and the treatments of vestibular dysfunction have gained momentum(Abolpour Moshizi et al., 2022; Purohit, 2023). Furthermore, studying the vestibular organs provides references for studying cochlea such as hair cell regeneration(Gonzalez-Garrido et al., 2021; Taylor et al., 2018), hair cell planar cell polarity establishment(Jiang, Kindt, & Wu, 2017; Kindt et al., 2021) due to their similar hair cell function(Curthoys, Grant, Pastras, Fröhlich, & Brown, 2021), and developmental processes(Atkinson, Huarcaya Najarro, Sayyid, & Cheng, 2015; Groves, Zhang, & Fekete, 2013). However, it is worth noting that standard, efficient, and comprehensive vestibular function tests for mice remain elusive, which prevents the results from different studies from being compared and thus results in an obstacle in vestibular-related research.

Mice are one of the commonly used animal models in research. In mice, common approaches for assessing vestibular function include morphological evaluation, behavioral tests, vestibular evoked potentials, and Vestibulo-Ocular Reflexs (VORs). Morphological evaluation(Balmer & Trussell, 2022; Nakayama, Helfert, Konrad, & Caspary, 1994) is relatively time-consuming, often requires sacrificing the mice, and is therefore not suitable for studies requiring significant amounts of animals such as drug screening(Ou, Santos, Raible, Simon, & Rubel, 2010) and longitudinal studies(Dall’Ara et al., 2016). Due to the involvement of multiple neural pathways(Trune & Lim, 1983), the commonly used behavioral tests, such as circling and head-tossing are not vestibular-specific. More specific behaviors, such as cycling and tilt, often rely on observer judgments and thus lead to subjective results(Dror, Taiber, Sela, Handzel, & Avraham, 2020; Hardisty-Hughes, Parker, & Brown, 2010; Zhao et al., 2021). While the vestibular evoked myogenic potentials (VEMPs) are a valuable clinical tool for assessing vestibular function, their application in mice is limited due to technical difficulties(Negishi-Oshino, Ohgami, He, Li, et al., 2019; Negishi-Oshino, Ohgami, He, Ohgami, et al., 2019). The short latency linear vestibular sensory evoked potentials (VsEPs) have been a commonly used method to analyze otolith function in mice(Gaines & Jones, 2013; Morley, Lysakowski, Vijayakumar, Menapace, & Jones, 2017; Vijayakumar et al., 2017). However, Jones. et al(Jones & Lee, 2021). mentioned that VsEPs can be interfered with by auditory brainstem response (ABR) evoked by the acoustic signs generated from the mechanical shaker during acceleration pulse production. Also, VsEP cannot be used to evaluate the semicircular canal deficits. VOR is a common test in clinics(Starkov, Strupp, Pleshkov, Kingma, & van de Berg, 2021) and has also been a standard vestibular function test in studies using monkeys as subjects(Cohen, Suzuki, & Raphan, 1983; Kushiro et al., 2002; Minor & Goldberg, 1991). VOR can be classified into angular VOR (aVOR) induced by the rotational accelerations and primarily driven by signals from horizontal semicircular canal cristae(HSCC)(Fetter, 2007; Yang et al., 2020), and translational VOR (tVOR), such as linear VOR (LVOR) and off-vertical axis rotation (OVAR), can provide otolith stimulus(Maruta, Simpson, Raphan, & Cohen, 2001; Simon et al., 2021b). LVOR is too small to provide reliable estimates of otolith function(Nist-Lund et al., 2019), and OVAR is easy to integrate with aVOR into one machine. Therefore, the combination of aVOR and OVAR can form the basis of a complete functional test for the peripheral vestibular organ. The commonly used techniques for recording VOR include video oculography (VOG), search coil, and electro-oculography (EOG), with VOG being the most commonly used recording method due to its high efficiency and non-invasive characteristics(Yang et al., 2020). Various systems based on the VOR have been developed for vestibular function test of mice(Harada et al., 2021; Khan, Brichta, & Migliaccio, 2022; Khan, Della Santina, & Migliaccio, 2019; Ono et al., 2020; Takimoto et al., 2018; Vijayakumar, Jones, Jones, Tian, & Johnson, 2019). Armstrong. et al.(Armstrong et al., 2015) implemented a vestibular function testing system in mice using hVOR and pOVAR(Shimizu et al., 2015) and evaluated the vestibular function of Caspase-3 mutant mice using this system. However, the system was not validated with common vestibular deficit mouse models. Additionally, no commercially available system made it challenging for other research laboratories to replicate them due to difficulties in integrating various components. For hearing tests on mice, TDT is a standard equipment, which allows a regular lab to perform hearing tests daily. There is no standard equipment for vestibular function tests on mice. This test has been very specialized with complicated equipment systems. And thus only a few labs can perform and other labs will have to send their mice to those labs.

Developed from an aVOR system(Yang et al., 2019), we implemented a comprehensive mouse vestibular peripheral functional evaluation system, integrating both aVOR and OVAR tests in a compact commercially available instrument. UVL can also be identified with a newly developed velocity steps stimulus. To validate the efficacy, we systematically tested the vestibular functions of the typical mouse models in vestibular research and demonstrated their accuracy and comprehensiveness in evaluating the common vestibular dysfunctions. The vestibular deficit mouse models are: 1) The dose-dependent vestibulotoxic drugs model was created by intraperitoneal injection of 3,3’-iminodiproprionitrile (IDPN) at three different doses (0, 2 and 4 mg/g). 2) The unilateral vestibular lesion (UVL) model was established by injecting the ototoxic drug gentamicin through semicircular canals. 3) Critical MET-related mutant mice (*Cdh23^v2J/v2J^*, *TMC1^-/-^*) were utilized in this study. 4) Vestibulo-specific mutant mice (*Zpld1^-/-^* for semicircular canal dysfunction, *Otop1^tlt/tlt^* for otolith deficit) were also tested with the system. Expected results were achieved, which demonstrated its potential as a standard system for mouse vestibular function evaluation.

## Materials and Methods

### The vestibular function test system

A schematic overview of the binocular VOG-based VOR test system (Shenzhen Giant Tek Co., Ltd) is provided in Figure 1. A microcontroller unit (MCU) controls a servo motor system to generate rotational motion of various profiles as the vestibular stimulus. The MCU interfaces with an LCD touchscreen for inputting the test parameters for those profiles, which are listed in the table in Figure 1A. The MCU also controls the video camera for recording the eye movement of the animal. The recorded video can be downloaded to a PC or edge computing device via a WIFI connection for further processing. A similar system has been previously described(Yang et al., 2019). The eye movement video recording and the data process are automatic. No human interaction is required. The video recording is synchronized with rotational stimulus. The stimulus parameters (for aVOR, frequencies, and speeds), sets up the test profiles. The eye movement during each different test profile is recorded in a separate video file. To accommodate the new OVAR test, the equipment system is hinged to a 50*50*2 cm aluminum base, which holds a motorized push rod to raise the back of the system and create a pitch of set angles.

**Figure 1.**
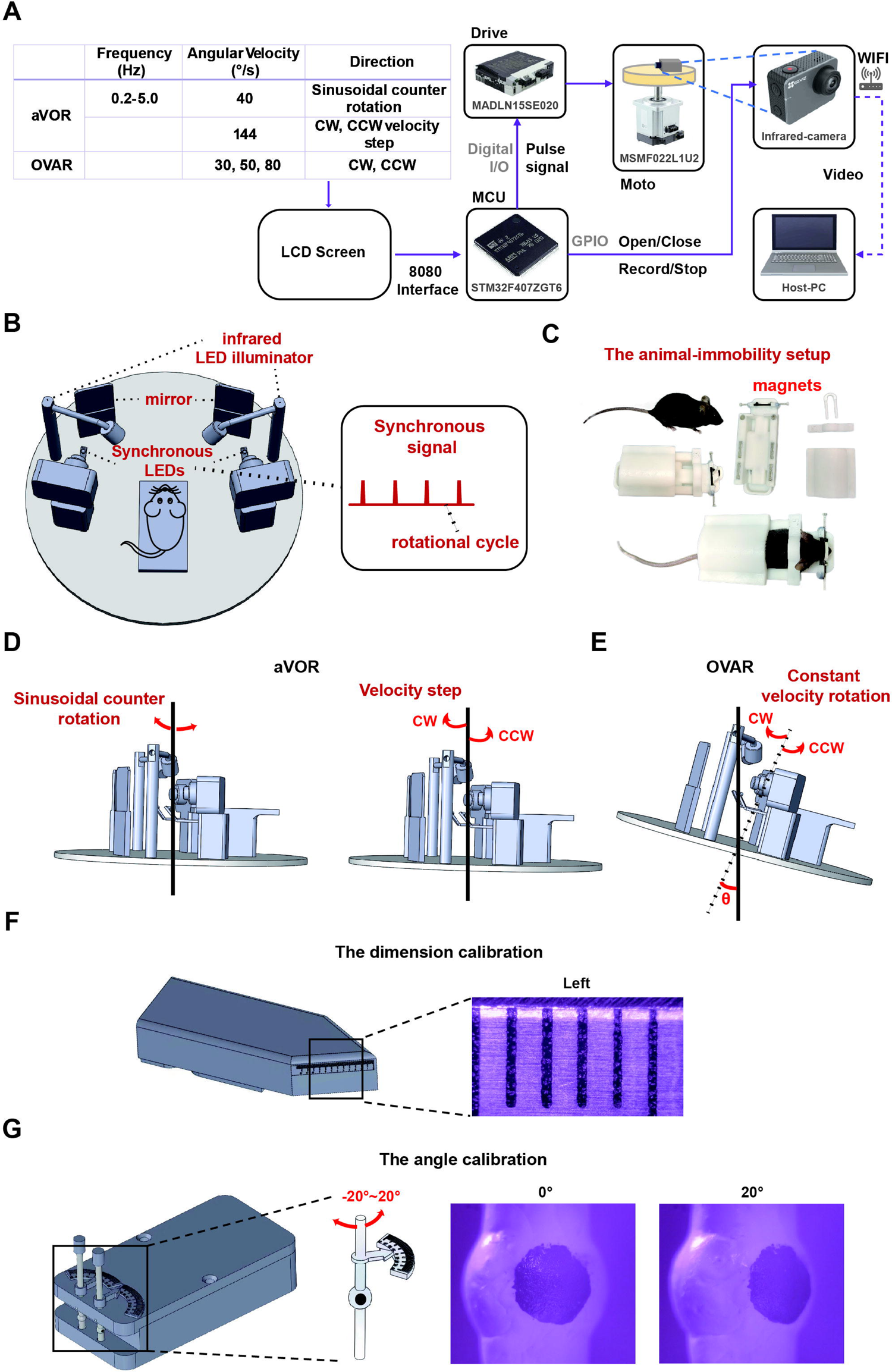
The mouse vestibular function evaluation system used in the studies. (A) The rotation modes of aVOR and OVAR used in the studies and the hardware architecture diagram of the VOR detection system. (B) Overview of the components of the system. (C) The noninvasive animal-immobility setup. In the setup, mice are head-fixed at ∼30° nose-down position to maintain the horizontal semicircular canals parallel to the yaw plane. (D) Scheme of rotation paradigm for the angular horizontal vestibulo-ocular reflex (aVOR) stimulus, a sinusoidal counter rotation, and velocity step stimulus in the horizontal plane. (E) Schematic view of apparatus for Off Vertical Axis Rotation (OVAR) stimulus, clockwise (CW) and counter-clockwise rotations (CCW) in yaw plane of 17° or 30° tilted platform at a rotation of speed of 30, 50, 80°/s. θ indicates the tilt angle of the rotation axis during OVAR. (F) The dimensions calibration of the eye movement recording system. (G) The angle calibration of the eye movement recording system.

Figure 1B shows that the platform holds the core of the test system: the animal and eye tracking system. The servo motor rotates the platform, which delivers this vestibular stimulus to the animal. Eye movement of the animal was recorded by two infrared-red motion cameras (EZVIZ S6, China) of 1920*1080 in resolution and 120 frames per second in speed. Illumination of the video recording was provided by two infrared LEDs, one for each eye. Another pair of LEDs flash synchronously with the rotational platform and marks the beginning and end of each rotational cycle, to calculate the phase delay of the eye movement to the sinusoidal rotational stimulus. The mouse was restrained in a custom-built holder adapted to its weight (Figure 1C) and then attached to a platform driven by the system. The holder allows for fixing the animal without anesthesia, which simplifies the preparation. The magnetic holder speeds up the fixing to about 30 s per animal (see the YouTube videos). Pilocarpine eye drops (2 %) were applied to the eyes to maintain pupil constriction in darkness.

### The test modes for evaluating vestibular functions

Two commonly used vestibular testing modes, angular Vestibulo-Ocular Reflex (aVOR) (Figure 1D) and Off Vertical Axis Rotation (OVAR) (Figure 1E), were integrated into this system.

The aVOR is generally used to assess HSCC function(Yang et al., 2020). Two types of stimuli, horizontal sinusoidal oscillation, and velocity step, were implemented to measure the HSCC function and to identify the side of unilateral vestibular lesion (UVL), respectively. During the HSCC test, the animal was stimulated by the sinusoidal oscillation of the platform at frequencies of 0.2, 0.5, 0.8, 1.6, 3.2, and 5.0 Hz, with peak velocities of 40 °/s. During the UVL test, the velocity step or trapezoid (144 °⁄s) stimulus was employed to identify the lesion side. The eccentric platform was accelerated for one second, then at a constant velocity for 32 seconds, and finally the velocity decreased to zero within one second. Horizontal nystagmus was observed for a while as the platform was accelerated or decelerated, from which the vestibular time constant was calculated as described in the previous study(Simon et al., 2021a).

The OVAR test has been used to evaluate otolith organ function in mice(Fetter, 2007). During the OVAR test, the equipment was tilted at 17° or 30°, as shown in Figure 1E. The platform rotated at constant velocities of 30, 50, and 80°/s in clockwise (CW) or counter-clockwise (CCW) directions. The horizontal and vertical (mostly in this paper) eye movements during the test were analyzed to evaluate the otolith function.

### Eye rotation calibration

The dimension and angle calibrations used in this study are shown in Figures 1F and G. The zoom factors of the two cameras in the system were adjusted to ensure they were identical. The field of view of both cameras is adjusted to 6 mm. An angle calibration was established using an ocular globe of 3 mm in diameter, about the same size as the eyeball of a 20 g mouse(Schmucker & Schaeffel, 2004). The eyeball diameter can also be changed according to the size of the mouse in the eye-tracking software. During the calibration, the artificial eyeball is rotated at precise angles with a protractor and the image of the artificial eyeball at different angles is recorded. The angular calibration scale can be calculated based on the translational distance of the pupil in the image and the angular readout from the protractor. The standard equipment and test procedure, and these calibration tools ensured the repeatability between different systems, making it possible for comparing results from different labs.

### Eye Movement Analysis

The eye movement analysis is implemented in an embedded computer: Jetson Orin Nano with parallel processing, as shown in Figure 2A. The video file for each test session was read by a CPU thread. Frames of the video were decoded and sent to a GPU thread for recognizing the pupil using a pre-trained YOLO v5 network. After a pupil in each frame was identified, the cropped pupil was sent to another CPU thread, where the pupil was segmented from the background, and the resulting binary image was then fitted by an ellipse to calculate the pupil center position. The two CPU threads and the GPU thread worked in parallel to compute the results, so that all results can be attained just a few minutes after the final experimental session. Eye movement is calculated by tracing the pupil center in horizontal and vertical directions (Figure 2B, C). Slow and fast components of the eye movement are extracted for evaluating different vestibular deficits.

**Figure 2.**
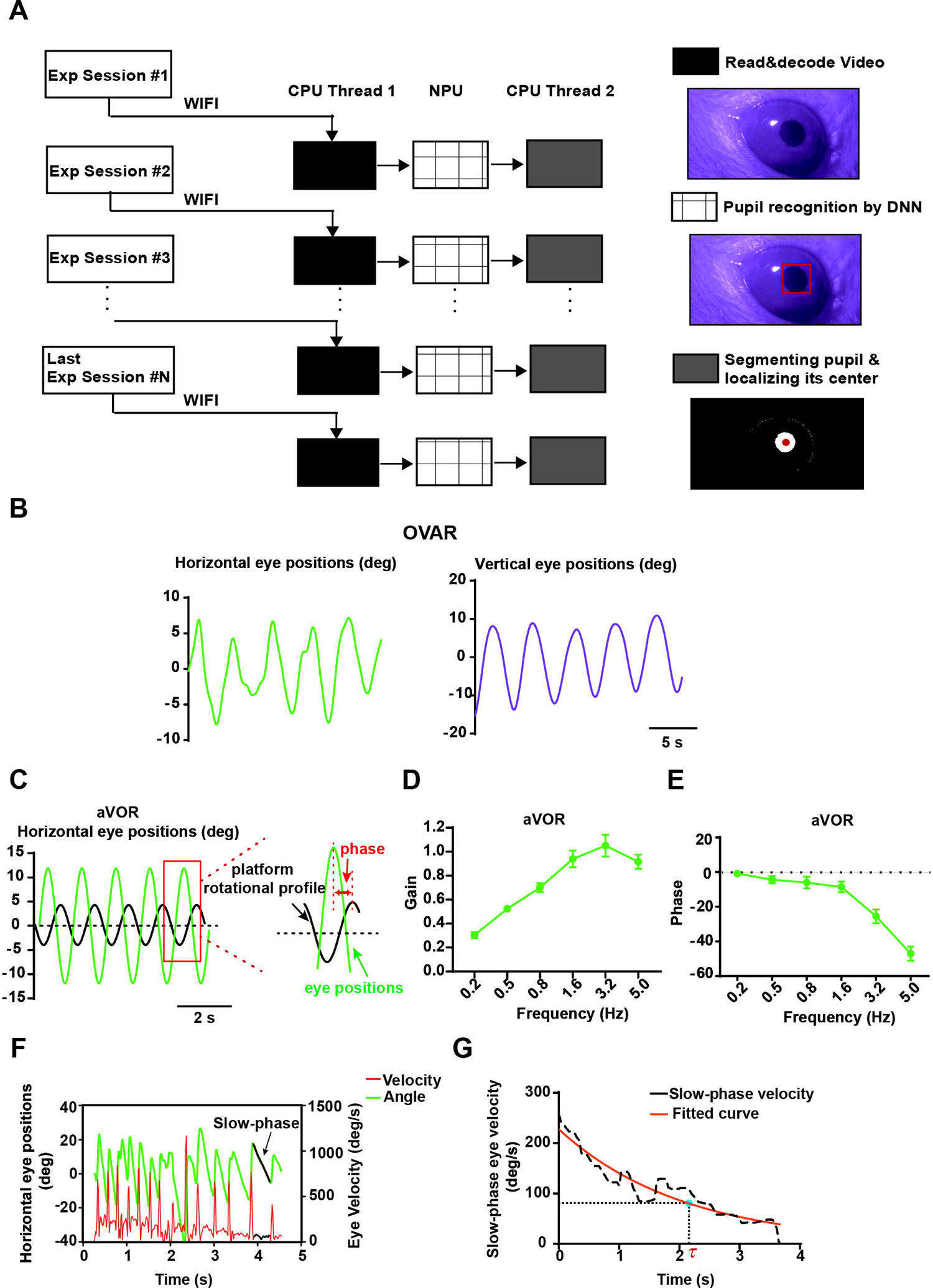
The workflow and outputs of the eye movement analysis. (A) The main steps are opening a YOLOv5 network, creating a project, selecting frames, labeling frames, and training a network. (B) The example outputs of measurable eye movements evoked by OVAR (80 °/s) include horizontal eye position and vertical eye positions. (C) The example outputs of aVOR responses (0.8 Hz) and the aVOR phase. Gain (D) and phase (E) for each testing frequency were computed from the aVOR waveforms. (F) The fast-slow alternating nystagmus and eye velocity were recorded at the beginning of the velocity step stimulus (The black arrow indicates the slow phase). (G) The horizontal slow phase velocity was fitted, and the vestibular time constant was calculated.

The eye movements evoked by OVAR include horizontal and vertical components (Figure 2B). The performance of the aVOR can be characterized by its gain (eye velocity/head velocity) and phase in the horizontal direction. Gain (Figure 2D) and phase (Figure 2E) for each testing frequency were computed from the aVOR waveforms using the methods described in detail by Stewart. et al(Stewart et al., 2016). The phase of aVOR represents the temporal shift between the eye and table rotations, expressed in degrees (Figure 2C).

The angular velocity step at 144 °/s stimulus was employed to evaluate the response of the vestibular center to HSCC input. At the beginning and end of the stimulus, a fast-slow alternating nystagmus will appear. As shown in Figure 2F, the eye velocity includes fast-phase velocity and slow-phase velocity. To calculate the vestibular time constant, the fast-phase component is removed, and an exponential fit is applied to the slow-phase eye velocity, as shown in Figure 2G. According to the curve *(f(x) =a * e^b*X^)*, the slow phase velocity is fitted, and the vestibular time constant is calculated as *r= −1/b*. By calculating the time constants of the vestibular system in both CCW and CW directions, it can reveal the asymmetry of peripheral input. This can be used to assess UVL.

### Mouse models of peripheral vestibular deficits

To verify the efficacy of the system in vestibular function evaluation, we adopted commonly used vestibular deficit mouse models with genetic, vestibulotoxic, and surgical manipulation. Table 1 lists the details of these models. 6-8-week-old C57BL/6J mice including both genders were used for the investigation of vestibulotoxicity drugs. The mutant mice and their wild-type littermates were used at 6-8 weeks old. All surgical and experimental procedures were approved by the Animal Care and Ethics Committee of the Southern University of Science and Technology.

**Table 1.**
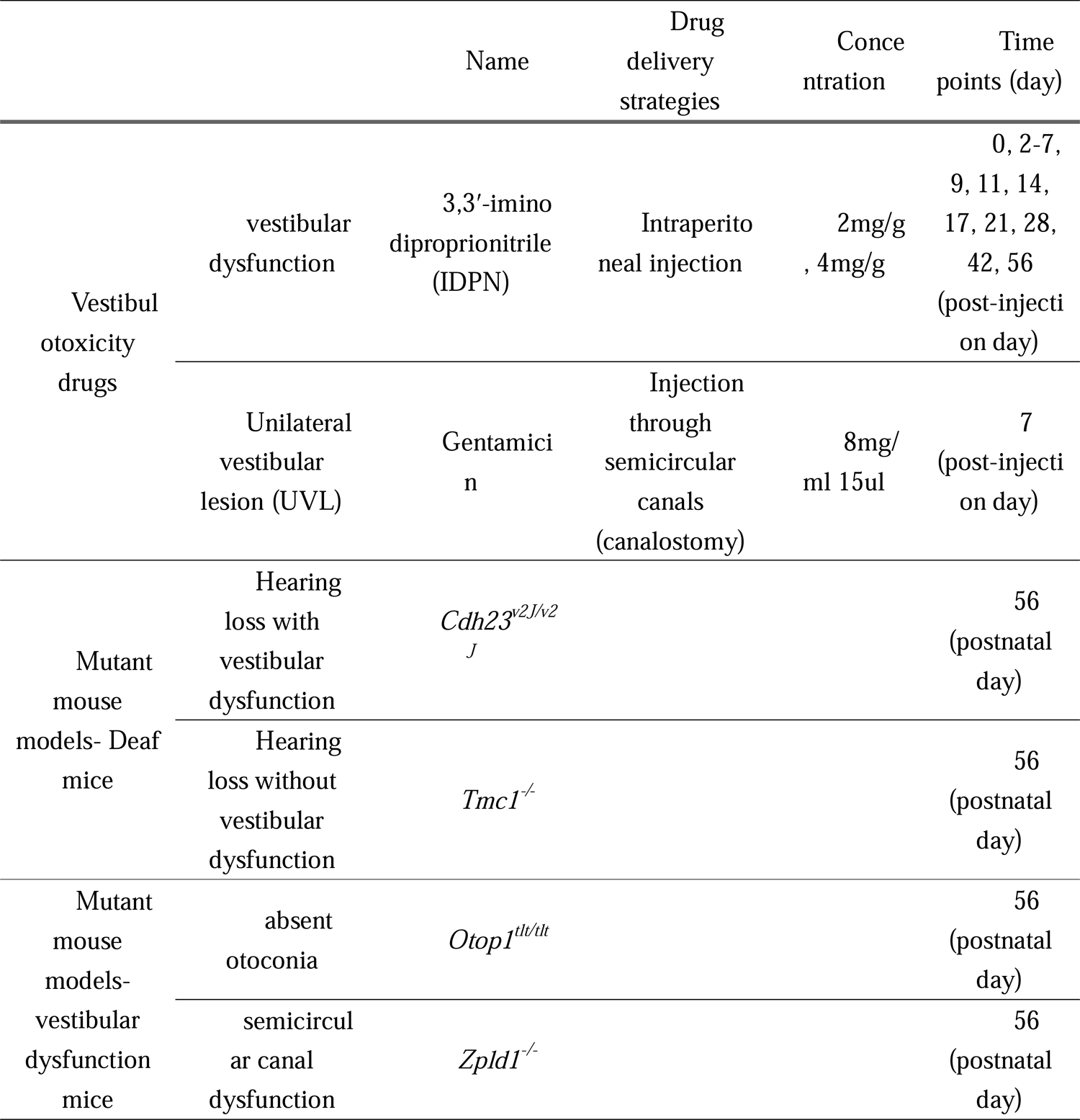

**Two mutant mouse models of mechano-electrical transduction deficits,** *Cdh23^v2J/v2J^*, and *TMC1^-/-^* were tested with the system. The Cdh23 protein is localized in the tip links of hair cell stereocilia and is proposed to regulate the activity of mechanically gated ion channels in mice(Siemens et al., 2004). The Tmc1 protein is a pore-forming component of mechanosensory transduction channels in hair cells(Kawashima et al., 2011). *Cdh23^v2J/v2J^*mice (The Jackson Laboratory, 002552) are characterized by circling, head-tossing, deafness, and hyperactivity. *TMC1^-/-^* mice were generated using Biocytogen Extreme Genome Editing System (EGE®) (Biocytogen, Beijing, China). *TMC1^-/-^*mice are deaf but with normal vestibular function due to the compensation of its closely related paralog *Tmc2*.

**Another two mutant mouse models with dysfunction at different vestibular organs,** *Otop1^tlt/tlt^* and *Zpld1^-/-^* were tested to demonstrate the specificity of the system. It has been reported that the zona pellucida-like domain protein (Zpld) is one major structural protein of the complex cupula structure(Dernedde et al., 2014). Moreover, Vijayakumar S. et al.(Vijayakumar et al., 2019) have reported that spontaneous mutations of the *Zpld1* gene in mice cause semicircular canal dysfunction but do not impair gravity receptor or hearing functions. Otop1 has been regarded as one important protein in regulating multiple aspects of otoconia and otolith development(Lundberg, Xu, Thiessen, & Kramer, 2015). The *Otop1^tlt/tlt^* mice carry single-point mutations that specifically impair the morphogenesis of otoconia without affecting other organs(Hurle et al., 2003). *Zpld1^-/-^* mouse model was generated using Biocytogen Extreme Genome Editing System (EGE®) (Biocytogen, Beijing, China). *Otop1^tlt/tlt^* mice (The Jackson Laboratory, 001104) are recessive mutations that affect balance by impairing otoconial morphogenesis without causing hearing loss(Hurle et al., 2003).

**Vestibulotoxic drug 3,3**′ **-iminodiproprionitrile** (IDPN, TCI, I0010, *d*=1.02 g/mL) was used to establish a vestibular dysfunction mouse model with quantified damage levels. IDPN, as a nitrile, is a neurotoxic and vestibulotoxic compound that has long been used to induce vestibular damage.(Llorens & Dememes, 1994; Llorens, Dememes, & Sans, 1993; Zeng et al., 2020). Each animal received a single intraperitoneal injection of IDPN at doses of 2, or 4 mg IDPN per gram of body weight, or saline (NS) as a control. The mouse vestibular function evaluation was performed before and after the drug exposure. During a period of 8 weeks, the three groups of animals were tested in 15 individual days, to show the progressive damage and recovery/compensation of the vestibular function.

**A unilateral vestibular lesion (UVL) mouse model** is established by delivering gentamicin (Macklin, G810322) with **canalostomy**, which is a surgical approach for local drug delivery into semicircular canals(Guo et al., 2018; Nakagawa et al., 2003). Briefly, a mouse was anesthetized by intraperitoneal injection of Dexmedetomidine Hydrochloride Injection (Dexdomitor^®^, 0.08 mg/kg) and Zoletil 50^®^ (32mg/kg). The surgical operation was performed on either the left or right ear. A post-auricular incision was made along the retroarticular groove, and the post-auricular region was prepared by shaving with a clipper and depilatory cream. The muscle covering the temporal bone was dissected using micro-forceps, revealing the posterior semicircular canal (PSCC) and HSCC. A small hole was created in the middle portion of the HSCC using a 26 G needle, and the presence of lymphatic fluid leakage confirmed the perforation of the bony wall. Subsequently, 15 μl of gentamicin (8 mg/ml) was injected into the HSCC via polyimide tubing (inner diameter 114.3 μm, outer diameter 139.7 μm) using a microinfusion pump (3 μl/min). After the injection, the hole was plugged with a small piece of muscle and covered with an adhesive agent. Finally, the skin incision was sutured.

### Auditory brainstem responses (ABRs)

The ABR measurements were conducted in a sound-proof chamber. The tone bursts from 4 to 32 kHz sound stimuli were delivered through a speaker (Tucker-Davis Technologies) placed in front of the animal 5 cm away from the vertex. ABRs were recorded via three needle electrodes: the active electrode was placed at the vertex of the skull; the reference electrode was placed at the mastoid area of the right ear, and the ground electrode was placed at the left hind leg. The response signals were digitized and acquired using BioSigRZ software (Tucker-Davis Technologies). The ABR threshold was defined as the lowest sound stimulus level that can evoke a clear and repeatable ABR waveform.

### Statistical Analysis

The “n” in the figures refers to the number of mice per group. All data are presented as the mean ± SEM. Data analysis was performed using GraphPad Prism 6 software. Student’s t-test was used for comparing two groups, while one- or two-way analysis of variance (ANOVA) tests were used for comparisons involving more than two groups. The detected difference was defined to be significant when *p* < 0.05.

## Results

### Quantification of IDPN-induced vestibular dysfunction and its progress

The IDPN-injected (NS, 2mg/g & 4 mg/g) mice were tested with different stimulus levels during a time course of 8 weeks (Figure 3A). There were 6 mice per group and three groups. We tested 18 mice per day with two VOR machines, one for aVOR and one for OVAR. Typical aVOR waveforms from the three groups of mice are shown in Figure 3B. Gain and phase for each testing frequency were computed from the recorded eye movement data. Typical waveforms of vertical eye movement in the OVAR test are shown in Figure 3C. Vertical eye movement amplitude in degrees was calculated from the waveforms.

**Figure 3.**
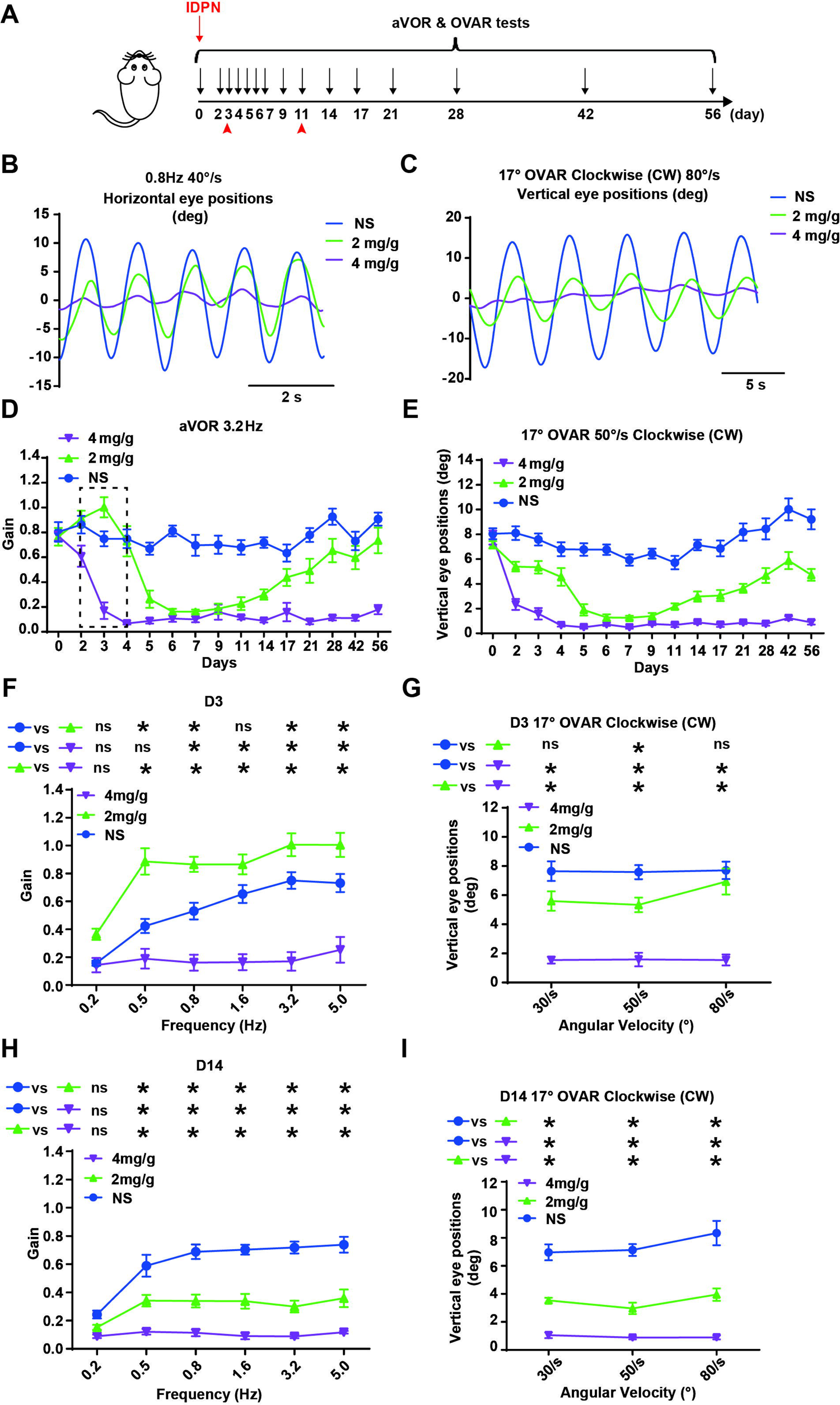
The vestibular dysfunction after IDPN injection. (A) Time course of vestibular system evaluation after IDPN injection. (B) Representative aVOR waveforms of the IDPN injected mice (NS, 2mg/g, and 4 mg/g) were induced by sinusoidal rotation of 0.8 Hz, peak velocity 40 °/s. (C) Vertical eye position waveforms of the IDPN injected mice (NS, 2mg/g, and 4 mg/g) were induced by OVAR stimulus (constant velocity of 80°/s in CW direction at the 17° tilted table). (D) The time-course characteristics of aVOR gain in the IDPN injected mice (NS, 2mg/g, and 4 mg/g, n=6 per group) at 3.2 Hz, 40 °/s. (E) The time-course characteristics of vertical eye positions in the IDPN injected mice (NS, 2mg/g, and 4 mg/g) at a constant velocity of 50°/s in CW direction at the 17° tilted table. (F) Gain of aVOR in the IDPN injected mice (NS, 2mg/g, and 4 mg/g) at 3 days (D 3). (H) Gain of aVOR in the IDPN injected mice at D 14. Amplitudes of vertical eye positions in the IDPN injected mice (NS, 2mg/g, and 4 mg/g) at 30, 50, and 80 °/s CW OVAR stimulus at D3 (G) and D14 (I). All data presented in mean±SEM, two-way ANOVA, * *p* < 0.05.

The progress of aVOR gain for the three groups under a rotational stimulus of 3.2 Hz and 40 degree/s was shown in Figure 3D (The gain at other frequencies, 0.2 Hz, 0.5 Hz, 0.8 Hz, 1.6 Hz, 5.0 Hz were shown in supplemental Figure S1A-E. All showed a similar pattern.). After IDPN administration, the aVOR gain of the 2 mg/g group shows a sharp decline on day 5 and a slow recovery from day 11. The aVOR gain of the 4 mg/g group showed an early sharp decline on day 3 and stayed at virtually zero thereafter. A similar pattern was shown in Figure 3E for the vertical movement amplitude in the OVAR test of CW-50 °/s stimulus (The amplitude under another stimulus was shown in supplemental Figure S1F-J. All showed a similar pattern.). Test results on two individual days (D3 and D14) were shown in Figure 3F and 3H for the aVOR tests, Figure 3G and 3I for the OVAR tests. The results, either from aVOR or OVAR test, demonstrated a dose-dependent decrease except for the aVOR responses at day 3. aVOR gain of the 2mg/g group increased at day 3 compared to the NS group under all stimulus modes. In prior work, we also found the aVOR gain of 2 mg/g group increase at an early stage of post-injection(Yang et al., 2019). These dose-dependent results and the progress pattern demonstrated the quantification capability of the test system to the vestibular function.

### Different vestibular phenotypes in MET-related mutant mice (*Cdh23^v2J/v2J^*and *TMC1^-/-^*)

Both adult *Cdh23^v2J/v2J^* and *TMC1^-/-^*mice are profoundly deaf (Figure 4A, B). The results of *Cdh23^v2J/v2J^* mice are shown in Figure 4C-G. *Cdh23^v2J/v2J^* mice have no obvious and recordable eye movements under all aVOR and OVAR stimuli (Figure 4C, F), but have spontaneous nystagmus (data not shown). Data shown in Figure 4D and 4E summarize the gains and phases of the aVOR responses for the *Cdh23^v2J/v2J^*mice and their wild-type littermates. While the WT group exhibited high gain compensatory responses at all the frequencies, *Cdh23^v2J/v2J^* mice exhibited virtually zero gain responses at the same frequencies, and therefore the phase of *Cdh23^v2J/v2J^* mice cannot be determined. As for the OVAR response, *Cdh23^v2J/v2J^* mice exhibited virtually no eye movement responses to head rotation (Figure 4G). The OVAR responses under 30° tilted table were similar to those of 17° (Figure S2A).

**Figure 4.**
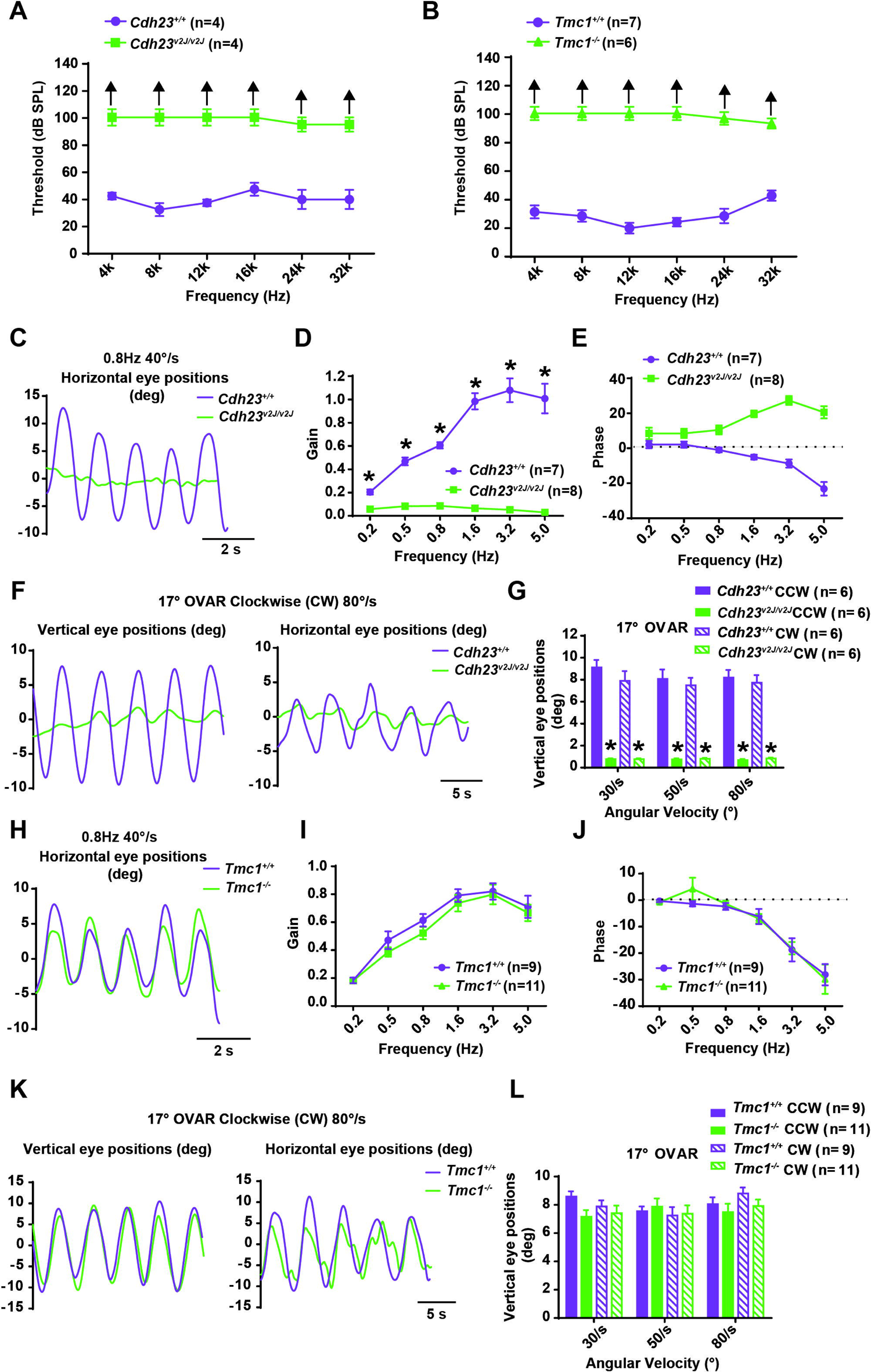
aVOR and OVAR responses in MET-related mutant mice (*Cdh23^v2J/v2J^* and *TMC1^-/-^*). The mean ABR thresholds of *Cdh23^v2J/v2J^* mice (A) and *TMC1^-/-^* mice (B) indicate that both of mice have severe hearing loss at all stimuli. (C) aVOR waveforms in response to 0.8 Hz 40 °/s stimulation of *Cdh23^v2J/v2J^* mice. Gain (D) and phase (E) of aVOR are plotted as functions of head rotation frequency for the WT group (purple) and *Cdh23^v2J/v2J^* mice (green). *Cdh23^v2J/v2J^* mice exhibited virtually zero gain responses and therefore the phase cannot be determined. (F) OVAR waveforms (horizontal and vertical eye positions) in response to constant velocity of 80°/s in CW direction at the 17° tilted table stimulation for *Cdh23^v2J/v2J^*mice. (G) Amplitudes of vertical eye positions in *Cdh23^v2J/v2J^*mice at 30, 50, 80 °/s OVAR stimulus. (H) aVOR waveforms in response to 0.8 Hz 40 °/s stimulation of the WT (purple) and *TMC1^-/-^* (green) mice. Gain (I) and phase (J) of aVOR for WT group and *TMC1^-/-^* mice. (K) OVAR waveforms (horizontal and vertical eye positions) in response to constant velocity of 80°/s in CW direction at the 17° tilted table stimulation for *TMC1^-/-^* mice. (L) Amplitudes of vertical eye positions in *TMC1^-/-^* mice at 30, 50, 80 °/s OVAR stimulus. All data presented in mean±SEM, unpaired t-test, * *p* < 0.05.

The results of *TMC1^-/-^* mice are shown in Figure 4H-L. *TMC1^-/-^*mice exhibited compensatory aVOR and OVAR that were similar to the responses of the WT mouse (Figure 4H, K). Gains of *TMC1^-/-^* mice were not significantly different from those of the WT group at all frequencies tested (Figure 4I). The phases were nearly identical to those of the WT group (Figure 4J). Similar OVAR responses were seen in WT and *TMC1^-/-^* mice across all stimulus modes (Figure 4L). The results of 30° OVAR were consistent with that of 17° in *TMC1^-/-^*mice (Figure S2B). These data demonstrated the capability of the test system in screening for the genetic mutants affecting vestibular function.

### Functional evaluation of semicircular canals and otolith organs respectively

The results of *Zpld1^-/-^* mice are shown in Figure 5A-E. *Zpld1^-/-^*mice have decreased aVOR response (Figure 5A) but normal OVAR response (Figure 5D). On average across frequencies of 0.2-1.6 Hz, aVOR gain was significantly lower in *Zpld1^-/-^* mice compared to WT mice (Figure 5B). There was no significant difference in aVOR phases between *Zpld1^-/-^* and WT mice under all stimulus modes (Figure 5C). Robust OVAR responses at all stimulus modes were seen in *Zpld1^-/-^* mice in response to head rotation. There was no significant difference in vertical eye movements between *Zpld1^-/-^* and WT mice at both 17° and 30° tilted table (Figure 5E, S2C).

**Figure 5.**
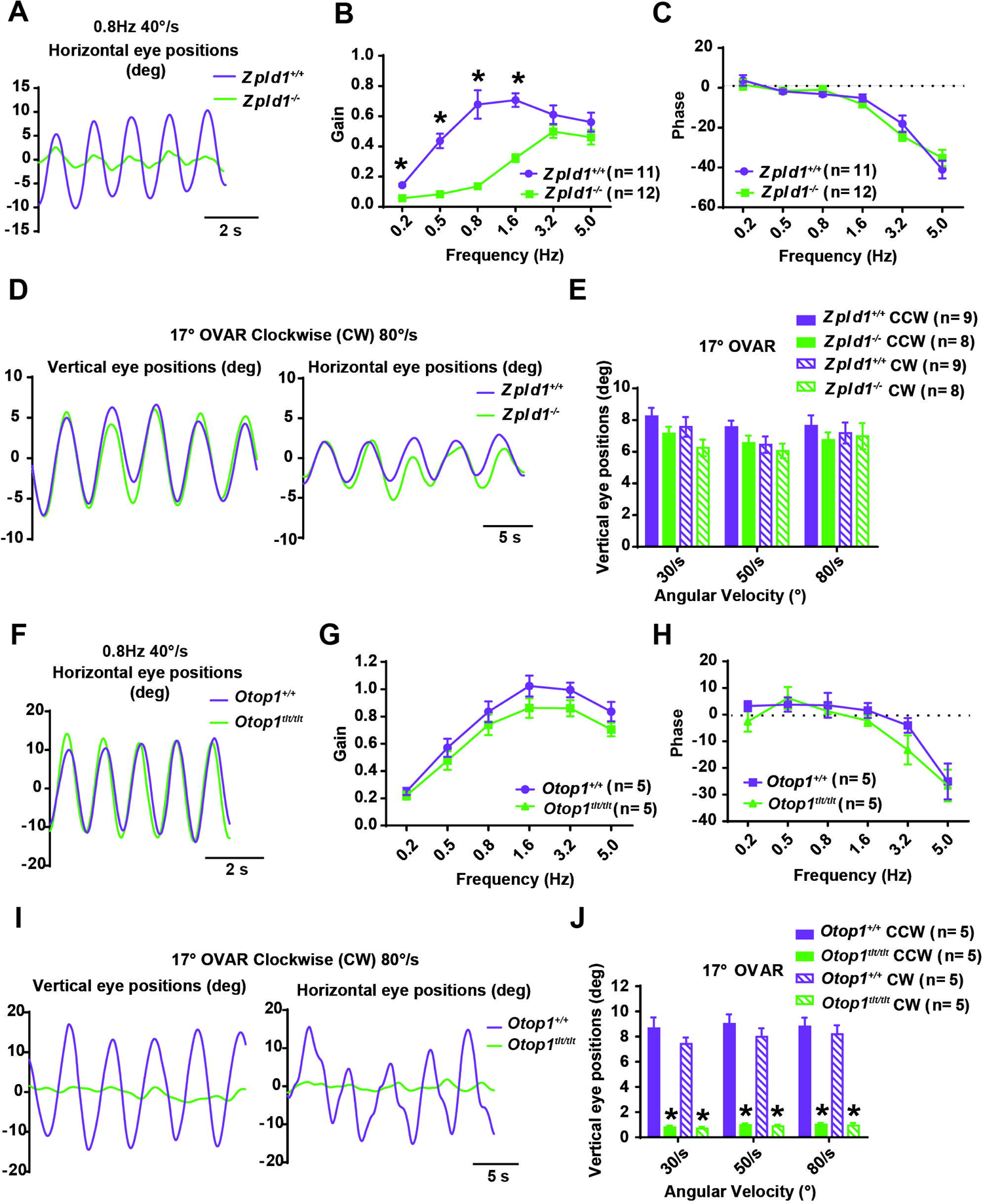
aVOR and OVAR responses in semicircular canal dysfunction mice (*Zpld1^-/-^*) and otoconia-deficient mice (*Otop1^tlt/tlt^*). (A) aVOR to sinusoidal head rotations with representative eye responses to 0.8 Hz head rotation for *Zpld1^-/-^* mice. aVOR gains (B) and phases (C) for the WT group (purple) and *Zpld1^-/-^* mice (green). (D) Representative eye position waveforms to constant velocity of 80°/s in CW direction at the 17° tilted table stimulation for *Zpld1^-/-^* mice. (E) Amplitudes of vertical eye positions of OVAR are shown for the WT group (purple) and *Zpld1^-/-^* mice (green). (F) aVOR to sinusoidal head rotations with representative eye responses to 0.8 Hz head rotation for *Otop1^tlt/tlt^*mice. aVOR gains (G) and phases (H) for the WT group (purple) and *Otop1^tlt/tlt^*mice (green). (I) Representative eye position measurements to the constant velocity of 80°/s in CW direction at the 17° tilted table stimulation for *Otop1^tlt/tlt^* mice. (J) Amplitudes of vertical eye positions in the WT group (purple) and *Otop1^tlt/tlt^* mice (green) at 30, 50, and 80 °/s OVAR stimulus. All data presented in mean±SEM, unpaired t-test, * *p* < 0.05.

The results of *Otop1^tlt/tlt^* mice are shown in Figure 5F-J. *Otop1^tlt/tlt^* mice exhibited normal aVOR responses (Figure 5F) but impaired OVAR responses (Figure 5I). Their aVOR gains (Figure 5G) and phases (Figure 5H) were not statistically different from those of the WT mice at all frequencies. The *Otop1^tlt/tlt^* mice exhibited virtually no eye movement responses to 17° and 30° tilted OVAR stimuli (Figure 5J, S2D). These results suggested that the test system could respectively assess the functions of the otolith system and semicircular canals.

### Identification of the lesion side on UVL mice

The VOR results of the unilateral vestibular lesion (UVL) mouse model are shown in Figure 6A-H. Typical examples of aVOR recordings from each group are shown in Figure 6A. Compared with the control group, aVOR gains in the UVL groups were significantly reduced at all tested frequencies (Figure 6B). However, there was no significant difference between the left and right eyes in the UVL groups. The phases of the UVL groups were generally not significantly different from those of the control group (Figure 6C). Figure 6D shows examples of OVAR responses from each group 7 days after the lesion. The OVAR responses in UVL groups were significantly impaired at all stimulus modes (Figure 6E, S2E-S2G) compared with the control group; however, there was also no significant difference between the left and right eyes in the UVL groups.

**Figure 6.**
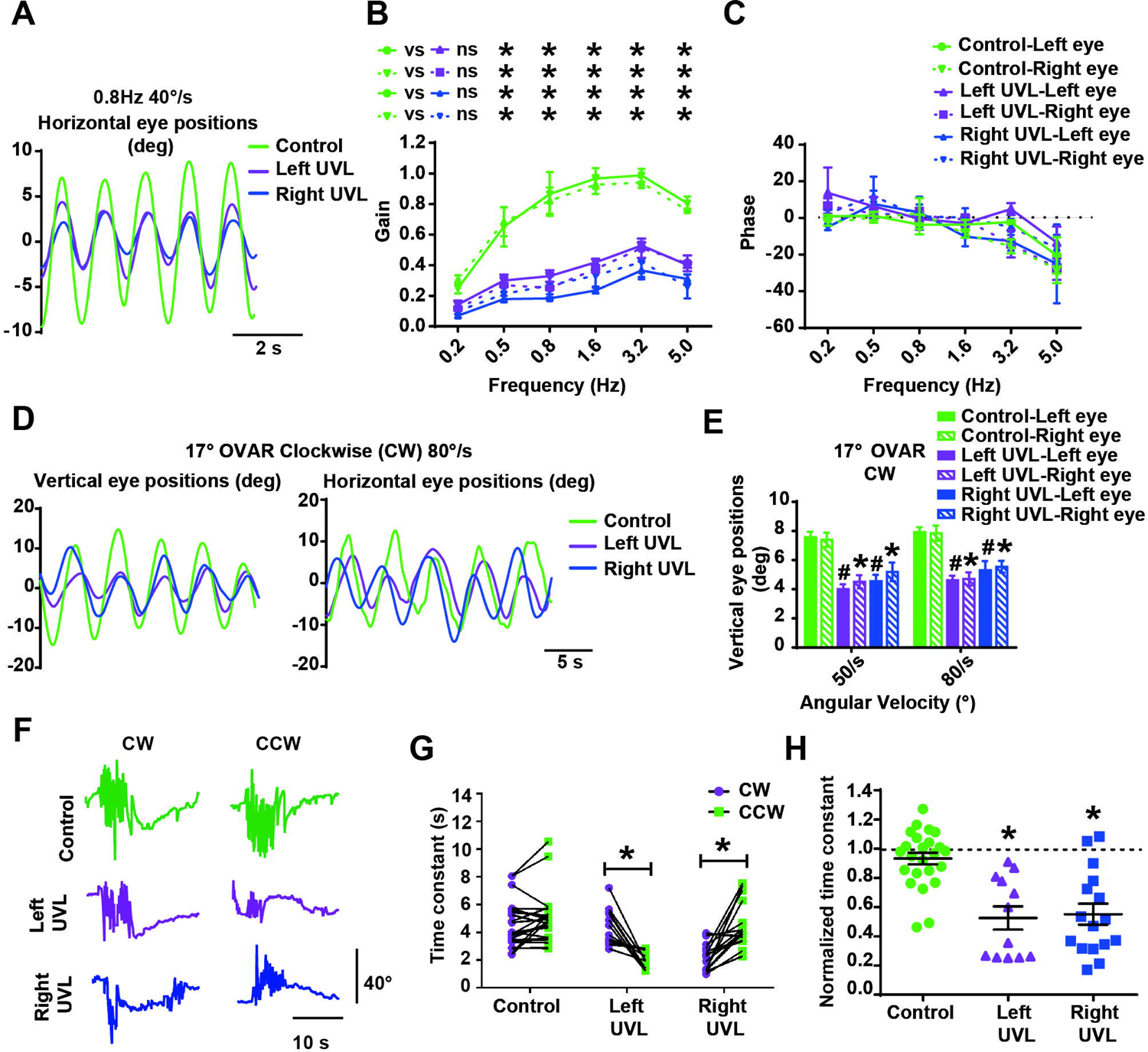
Impaired vestibular function in unilateral vestibular lesion (UVL) mice. (A)Typical aVOR waveforms of the control group (green), left UVL group (purple), and right UVL group (blue) were induced by sinusoidal rotation of 0.8 Hz, peak velocity 40°/s. Gain (B) and phase (C) of the left eye and right eye aVOR for the control group (green), left UVL group (purple), and right UVL group (blue). (D) Representative raw tracings of vertical and horizontal eye movements of the control group (green), left UVL group (purple), and right UVL group (blue) at 80 °/s OVAR stimulus. (E) Amplitudes of the left eye and right eye vertical eye positions in the control group (green), left UVL group (purple), and right UVL group (blue) at 17° tilted platform OVAR stimulus in CW direction, * *p* < 0.05 compared with Control-Right eye; # *p* < 0.05 compared with Control-Left eye. (F) Velocity step or trapezoid (144 °⁄s) stimulus was performed. Traces show horizontal eye position during velocity steps stimulus. (G) The time constant for the control group, left UVL group, and right UVL group in CCW and CW direction (paired t-test). (H) The normalized time constant for the control group (green), left UVL group (purple), and right UVL group (blue). The time constant at contralesional stimulation (CW for left UVL, CCW for right UVL) was taken as 1. All data presented in mean±SEM, two-way ANOVA, * *p* < 0.05.

During velocity steps stimulus, while the time constant obtained at the beginning of CW and CCW rotation was symmetric in the control group, that in UVL mice was asymmetrical and lateralized to the unaffected side rotation (CW in left UVL and CCW in right UVL) (Figure 6F, G). To simplify the comparison of the time constant between control and UVL mice, the time constant at unaffected side rotation (CW for left UVL, CCW for right UVL) was taken as 1 (Figure 6H). These results suggested that the contralesion and ipsilesion can be identified via the time constant.

## Discussion

In our study, we have successfully developed a vestibular function evaluation system, and its efficacy was demonstrated with comprehensive experiments using common vestibular-deficit animal models. The integrated aVOR and OVAR modes allowed us to assess the otolith organs and semicircular canals separately. UVL can be identified by quantifying the time constant during the velocity steps stimulus. The ease of use and reliability of the system allowed us to conduct hundreds of experiments within a span of two months to monitor the progress (deterioration and recovery) of the mouse vestibular functions after ototoxic drug IDPN treatment. This longitudinal study also helped discover the unexpected onset increase in aVOR gain for low-dose animals.

### Low-dose IDPN treatment resulted in increased aVOR gain at the early stage

Due to its convenience of use, our system allowed us to conduct long-term monitoring of vestibular function change following various treatments. As an example, during the IDPN toxicity experiment, we observed an increase in aVOR gain at the 2 mg/g dosage groups from Day 2 to Day 4 compared with the baseline before treatment. This phenomenon was also observed in our previous study(Yang et al., 2019). While the precise interpretation of this phenomenon remains elusive at present, there have been reported that hair cells may not be the primary target of IDPN(Seoane, Demêmes, & Llorens, 2001). Type I hair cells are high sensitive to IDPN, and their calyx junction dismantlement and synaptic uncoupling are early events in the mouse vestibular sensory epithelium after IDPN exposure(Greguske, Carreres-Pons, Cutillas, Boadas-Vaello, & Llorens, 2019; Maroto et al., 2023; Sedo-Cabezon, Jedynak, Boadas-Vaello, & Llorens, 2015). Research has indicated that exposure to IDPN may lead to alterations in the regulatory responses of vestibular afferent neurons to acetylcholine(Greguske et al., 2023). Acetylcholine is a neurotransmitter of the vestibular efferent nervous system, in which α9/10 nicotinic acetylcholine acts as a descending inhibitory neurotransmitter that can reduce the sensitivity of type II hair cells. We speculate that in the early events induced by IDPN, the regulatory capacity of the vestibular efferent system is diminished, leading to reduced inhibition of Type II hair cells. Consequently, there is a heightened response to rhythmic head oscillations, leading to the observed increase in aVOR at the 2 mg/g dosages during the early stages. Further studies are needed to test this hypothesis.

### VOR is a more reliable measure of vestibular function than behavioral tests

Behavioral tests are also a common measure of vestibular function for mice. During the *Zpld1^-/-^* experiment, only a few mice (two out of twelve) exhibited observable behavioral patterns such as circling behavior, and head bobbing. Vijayakumar and his colleagues also reported this highly variant behavioral pattern in the *Zpld1* spontaneous mutant mice(Vijayakumar et al., 2019). However, all *Zpld1^-/-^* mice consistently show reduced aVOR gain. This result suggests that VOR measurements may be a more reliable measure of vestibular function.

### The lesion side of UVL can be identified by VOR with an asymmetric time constant

Although VOR is thought not to be able to detect UVL, our data indicate that the time constant during the velocity step or trapezoid (144 °⁄s) stimulus provides a simple metric for detecting UVL. In cases of unilateral damage to the vestibular organ, there is a reduction or even silencing of resting firing frequency on the lesioned side, creating an imbalance between the normal resting firing rates on the contralesional and ipsilesional sides(Fetter, 2016). This imbalance mimics a constant rotation toward the contralesional side, resulting in asymmetric responses.

## Summary

In this paper, a mouse vestibular function test system was introduced and its efficacy was systematically tested with common peripheral vestibular deficit mouse models. The design of the animal holder together with the automatic test procedure allows researchers in a regular lab to perform the test daily. This system may provide standardized evaluation for mouse vestibular function and may also be useful for studying the development and regeneration of the inner ear. Evolving from an aVOR system(Yang et al., 2019), we extended the OVAR and velocity step or trapezoid (144 °⁄s) stimulus into our system for specific testing of the otolith organ. Other stimulus paradigms may be expanded in the future. Based on the system, mouse models of vestibular central deficit may also be evaluated by applying a new stimulus profile and extracting more eye movement features.

---

**Figure S1. The vestibular dysfunction after IDPN injection.** The time-course characteristics of aVOR gain in the IDPN injected mice (NS, 2mg/g and 4 mg/g) at 0.2 Hz (A), 0.5 Hz (B), 0.8 Hz (C), 1.6 Hz (D), and 5.0 Hz (E), 40 °/s. The time-course characteristics of vertical eye positions in the IDPN injected mice (NS, 2mg/g and 4 mg/g) at a constant velocity of 30 °/s (F), 50 °/s (G), 80 °/s (H) in CCW direction and 30 °/s (I), 80 °/s (J) in CW direction at the 17° tilted table.

**Figure S2.** Amplitudes of vertical eye positions in *Cdh23^v2J/v2J^*mice (A), *TMC1^-/-^* mice (B) *Zpld1^-/-^* (C) and *Otop1^tlt/tlt^* mice (D) at 30° tilted platform OVAR stimulus. Amplitudes of the left eye and right eye vertical eye positions in the control group (green), left UVL group (purple), and right UVL group (blue) at 17° tilted platform CCW direction (E), 30° tilted platform CCW direction (F) and CW direction (G) OVAR stimulus, * *p* < 0.05 compared with Control-Right eye; # *p* < 0.05 compared with Control-Left eye.

## Notes

### Competing Interest Statement

The VOR test system used in our study was developed by Giant Tek Ltd. Wenda Liu is an employee of Giant Tek, and Fangyi Chen is its technical advisor.

