## Supplementary figures and images for "An integrated system for comprehensive mouse peripheral vestibular function evaluation based on Vestibulo-ocular Reflex"

### Supplemental Figure 1

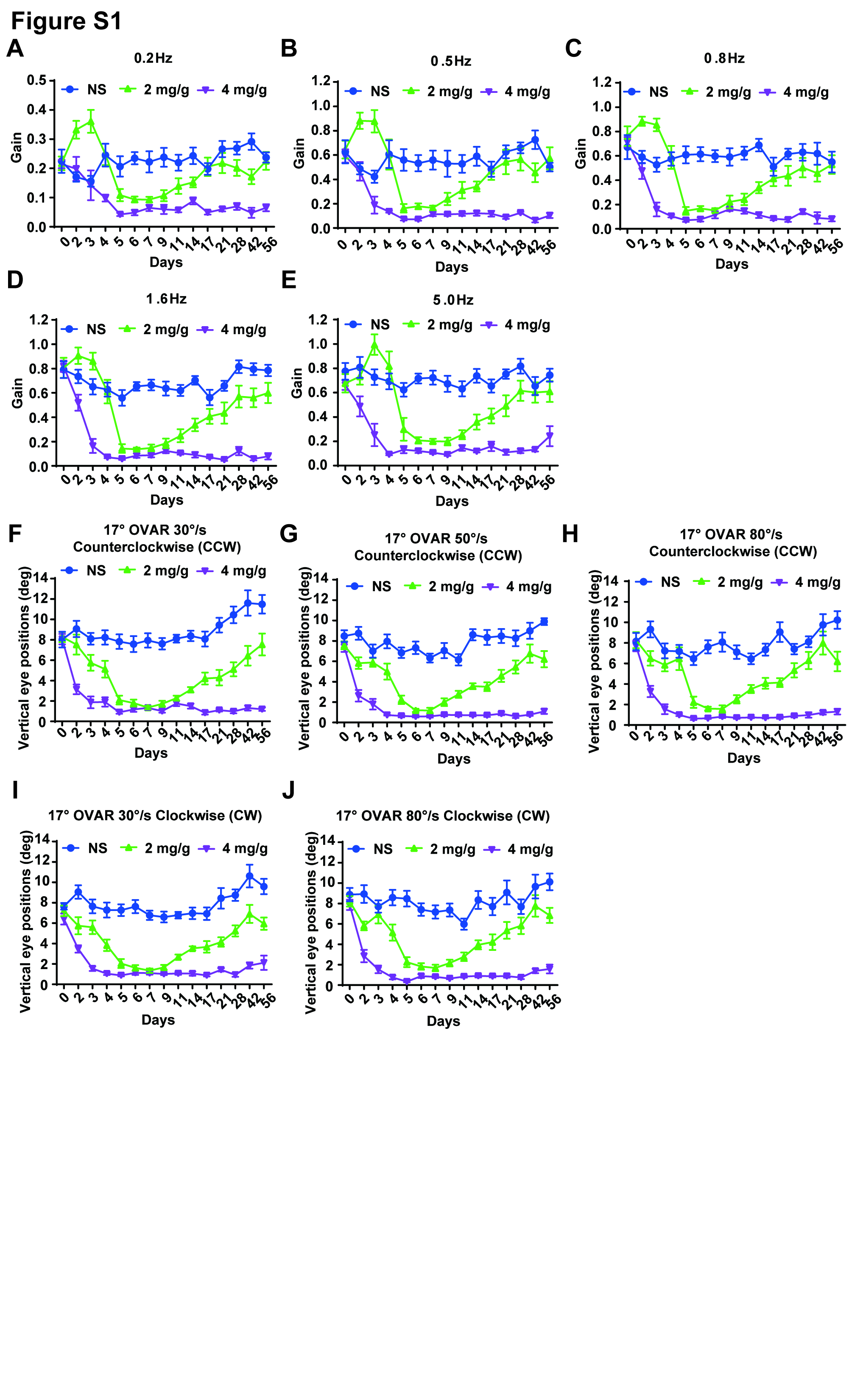

### Supplemental Figure 2

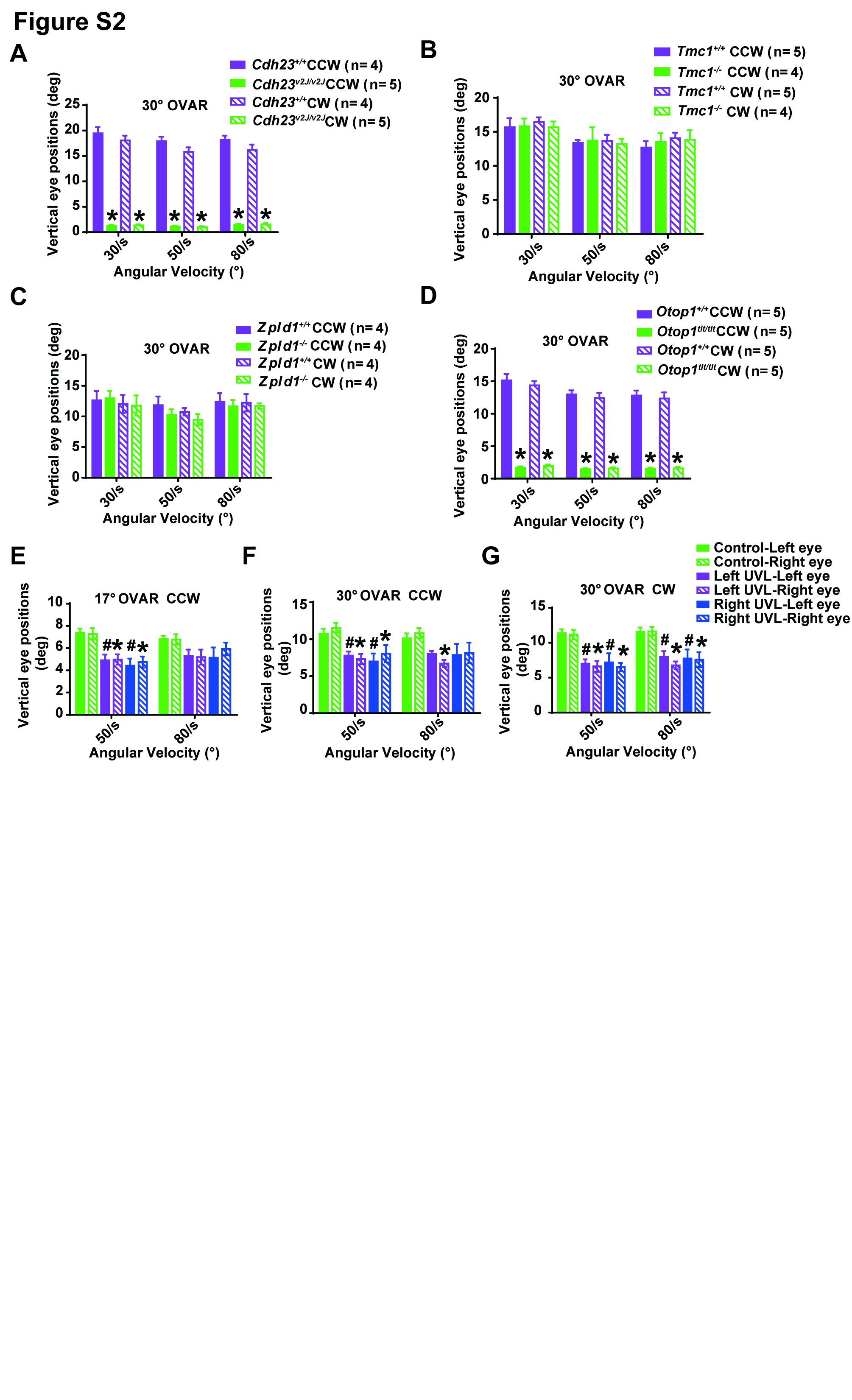
